# Discovery and shaping of decision strategies using adversarial stimuli in rats

**DOI:** 10.1101/2025.09.09.675127

**Authors:** Anqi Zhang, Anthony M Zador

**Affiliations:** Cold Spring Harbor Laboratory, Cold Spring Harbor NY; Cold Spring Harbor Laboratory School of Biological Sciences, Cold Spring Harbor NY; Department of Molecular and Cellular Biology, Harvard University, Cambridge MA

## Abstract

Animals typically learn to solve decision tasks in the laboratory through trial and error, rather than through explicit instruction of decision rules. The decision rule used by the animal may be difficult to read out directly from choice or accuracy data, when multiple decision rules are possible given the task design. Here, we demonstrate that in rats performing a visual decision task, probe stimuli can be used to gain information about decision strategy, and in our task revealed variation in the decision strategies used across rats. Further, we find that in a more general version of this task, rats use varying decision strategies that differ from the optimal ideal observer strategy, but respond to manipulations of the stimulus distribution by adjusting their behavioral strategy. Therefore, we show that informative probe stimuli can be used in both training and testing to confirm and shape behavioral strategy in a perceptual decision task.

## Introduction

During perceptual decisions, animals must process sensory inputs and produce the appropriate action that leads to reward. A crucial step toward understanding the neural basis of this sensorimotor transformation is to first understand how exactly animals use sensory information and past history to guide their decisions (Waskom et al, 2019). In simple decision-making tasks involving a binary decision and a reward contingency determined by the experimenter, the decision is primarily a function of the current trial’s stimulus. This relationship is typically fit well by a logistic function. However, though this function tells us that the stimulus is being used in the decision, it does not necessarily tell us the strategy that is used to make the decision. This becomes ambiguous when multiple strategies can be used to solve a particular problem.

Consider a task in which you are asked to report which of two numbers, each drawn from between 1-10, is greater. You will be most accurate if you consider both numbers and choose the greater. You will be less accurate, of course, if you simply base your decision on whether the first number is a low number (the second number is likely to be greater) or a high number (the second number is likely to be lesser). But, by completing many trials using the second strategy, you will still collect a great deal of reward, and because of the limited range of possible stimuli, your accuracy will still scale with the size of difference between the two numbers. How can the true strategy be measured in such a case? Here we use stimulus manipulations to directly probe decision strategy in rats performing a visual decision task.

Probe trials have been used in decision-making tasks to elucidate expectation of reward (Lak et al, 2014), and to disentangle latent learned associations from other processes such as exploration or information seeking (Drieu et al, 2025). In most instances, what differs in a probe trial is the availability of the reward or the context of the environment. Here we use specific points in the stimulus space as probe trials to infer animal strategy. We propose that “out-of-distribution” stimuli, to use vocabulary inspired by the machine learning field (Lee et al, 2018), and in particular those that are “adversarial” to a given candidate strategy – that is, those stimuli that would confuse strategy A but not an alternative strategy B – can be maximally informative about the use of strategy A vs B (vs C, and so on).

We applied this approach to a rodent visual decision task, and found that rats’ decision strategies were readily measurable from individual sessions. When we extended the space of possible stimuli, we found that rats’ decision strategies were again variable and differed from the optimal ideal observer strategy. This raised the question of whether we could further leverage adversarial stimuli to shape decision strategies. Indeed, we found that animal strategy could be shaped by exposure to a stimulus distribution that overemphasized these adversarial stimuli, suggesting that decision strategy is flexible and integrates experiences over a long time horizon.

## Results

We studied decision strategies using the visual discrimination task previously reported in Zhang and Zador (2023). We trained five rats to make a left or right choice report depending on the distribution of dots on a monitor placed directly behind a panel of three nosepokes. The visual stimulus consisted of white dots distributed across a black background, and rats were asked to report whether more dots were present in the upper or lower subregion of the stimulus. Each rat learned a particular contingency between the upper/lower distribution of the dots and the left/right binary choice reports. Rats were first trained on easy trials and after acquisition of the rule, were presented with more difficult stimuli (Figure 1a).

**Figure 1.**
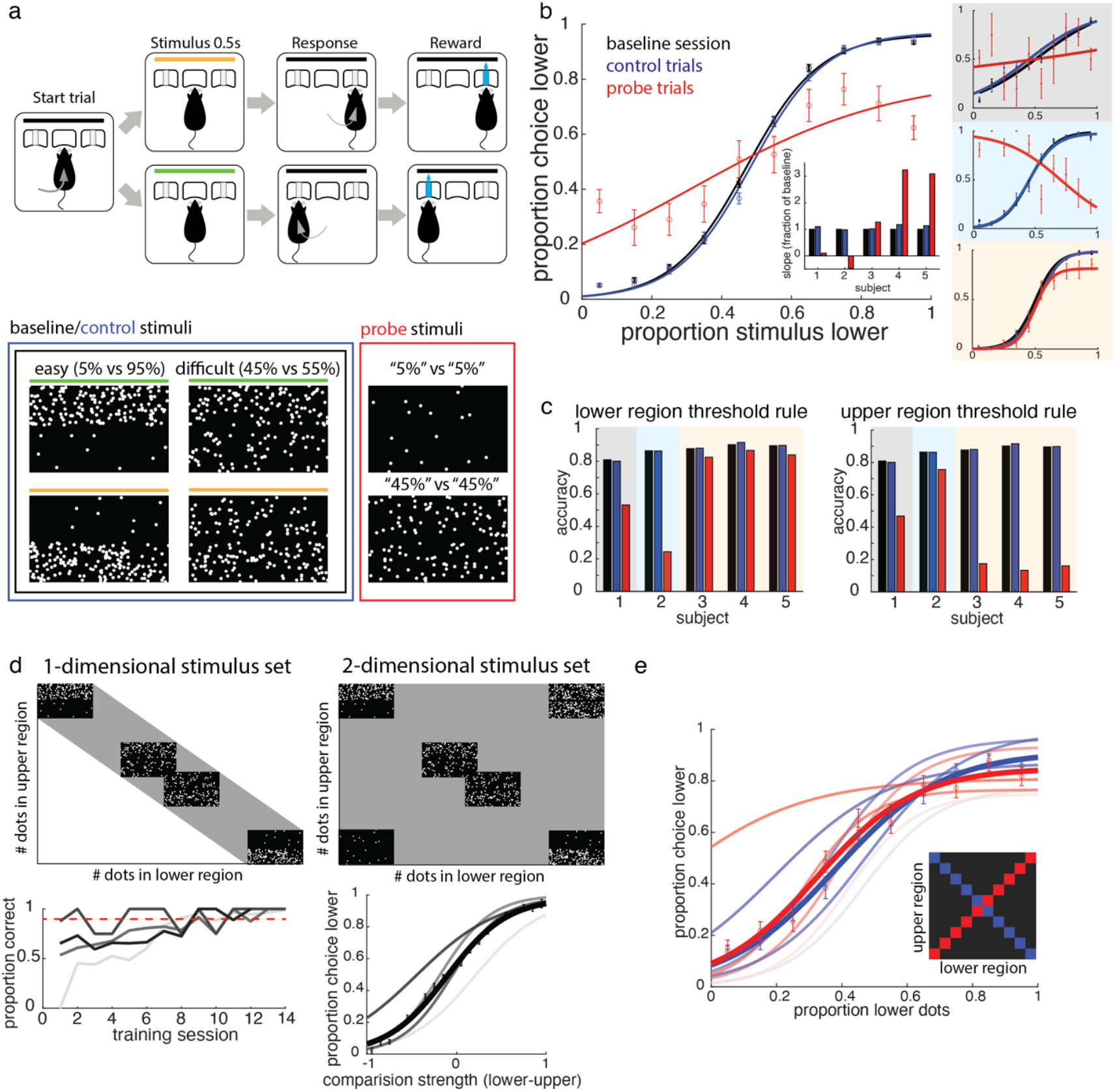
Probe stimuli are a window into underlying behavioral strategy. a) Visual decision task structure (previously reported in Zhang and Zador, 2023) and example stimuli. Animals are trained on baseline sessions (example stimuli in black box). After learning, they are tested on probe sessions, when probe (red) stimuli are delivered on 10% of trials and control (blue, same as baseline stimuli) are delivered on the remaining trials. b) Comparing psychometric curves between baseline sessions (black) and between control (blue) and probe (red) trials in probe sessions reveals differing strategies used by animals: comparison between upper and lower (grey shading), threshold applied to upper stimulus region (blue shading), and threshold on lower stimulus region (yellow shading). Inset: change in psychometric slope from baseline, for each of the three sets of trials. c) Decision rule preference is reflected in choice accuracy in response to probe stimuli, as measured according to each thresholding rule. A subject using the comparison rule (subject 1, in grey) would show chance accuracy by both of these decision rules. d) Modified task where stimulus covers a 2D space. Animals learn quickly over days, reaching 90% correct on easy trials in less than 14 days. Choice behavior is a function of comparison strength. e) Subsampling the trials in the 2D decision task to the baseline and probe trials analyzed in b) reveals animal strategy largely stays similar between the 1D and 2D tasks, and most animals follow a thresholding strategy. Inset: stimuli included in this analysis, with control trials in blue and probe trials in red.

Upon achieving expert performance on this task, we delivered probe stimuli on 5-10% of the trials, where the density of dots in the upper subregion of the stimulus was matched to the lower subregion (Figure 1a). We termed these sessions “probe sessions,” but in the majority of trials, typical stimuli were presented and rewarded according to the learned contingency. Probe trials were rewarded randomly to discourage learning of new associations. We continued to plot rat behavior as a function of the lower subregion stimulus strength, and observed that behavior on these probe trials varied across rats, with individual rats’ choice patterns aligning to one of three possible solutions to this task (Figure 1a, b).

In one rat, choice behavior became random and decoupled from stimulus strength, suggesting this rat’s good performance on control trials depended on a comparison between the content of the two stimulus subregions (Figure 1b, grey inset). When this comparison signal was abolished across probe trials, this rat’s performance approached random selection. In contrast, other rats’ performance showed markedly different patterns on the probe trials. In a second rat, the psychometric curve on probe trials flipped, but maintained a monotonic relationship with the stimulus strength (Figure 1b, blue inset). We realized that the decision rule applied by this rat was likely a threshold on the density of dots in the upper subregion of the stimulus. For example, when the lower subregion has a low density of dots (the leftmost point along the x axis on the blue curve), on control trials, the upper subregion has a high density of dots. In the probe trials, however, when the two subregions are equal in dot density, the corresponding datapoint where the upper subregion’s dot density is high is when the lower subregion’s dot density is also high (the rightmost point along the x axis on the red curve). We see that the rat’s choice profile at these two points is comparable, and as such, the rat’s performance on probe trials is consistent with a strategy of applying a threshold to the stimulus strength in the upper subregion of the screen. Finally, three of five rats demonstrated performance on probe trials that was very similar to their performance on control trials (Figure 1b, orange inset). By the same reasoning as for the second rat above, we inferred that these rats primarily use the lower subregion of the stimulus to drive their decision. As such, in probe trials, the animals still display the same choice behavior as on control trials when plotted against the lower subregion stimulus strength. We confirmed these interpretations by measuring how accurate each animal’s performance was in each condition according to either a lower or upper subregion thresholding rule (Figure 1c). Notably, we found that there was no change in performance on control trials within the probe session, compared to performance in baseline sessions just prior to the probe sessions. This reassured us that the delivery of probe stimuli on a small subset of trials did not perturb the animals’ performance on the remainder of the trials on that session.

We questioned whether the variability of decision rules used by different animals, and in particular, the reliance of most animals on a single subregion of the stimulus, was a byproduct of our simple one-dimensional stimulus design (Figure 1d). This stimulus set was designed to always present the same number of individual dots across all frames and all stimulus conditions, and as such, the upper and lower subregions of the stimulus provided complementary and therefore redundant information. We were surprised to note that the strategies employed by four out of five rats seemed to take advantage of this correlation. We considered whether such a strategy (biased use of one subregion) would continue to be selected if rats were trained on the full two-dimensional stimulus space containing all possible combinations of dot densities in the two subregions. Thus, we adapted our existing task to deliver stimuli uniformly selected from this space (Figure 1d). We trained 4 rats on this version of the task and found that they were able to readily learn this task within 14 sessions, which is comparable to the learning curves previously reported for the original task (Zhang and Zador, 2023). Performance at the end of learning was well-fit by a logistic function of the comparison strength between the two subregions. However, when we again subsampled the stimulus conditions used in our probe versus control trials in the original task, we found that the psychometric curves were nearly indistinguishable when plotted against the dot density in the lower subregion of the stimulus (Figure 1e), just like for rats 3, 4, and 5 in Figure 1b, indicating again that these rats showed a predominant reliance on the information in that part of the stimulus.

How can we reconcile the global behavior as seemingly being a function of the comparison strength, with performance on these subsets of trials demonstrating that rats can make decisions even at zero comparison strength? The reason is that when we uniformly sample the full two-dimensional stimulus space in Figure 1d, there remains a large degree of overlap between comparison strength and the density in each subregion. When we visualize decision rules as decision boundaries in the 2D stimulus space (Figure 2a), it becomes evident that the comparison strategy and single-region thresholding strategy agree in the majority of stimulus cases (black stimulus regions). Notably, the performance of an ideal observer on stimuli in the magenta stimulus regions, and therefore across the full stimulus space, will differ depending on which of the two rules is used. However, the introduction of sensory noise blurs this distinction. Given a fixed width Gaussian for sensory noise, choices made by a lower-thresholding strategy can be replotted as a function of the comparison strength between regions and fitted well by a logistic function (Figure 2b). The imperfect correlation between these two strategies means that the same choice profile forms a monotonic relationship with either stimulus parameter (single region thresholding or comparison strength), and thus the presence of a monotonic relationship is not sufficient to conclude the use of that stimulus parameter in driving those choices.

**Figure 2.**
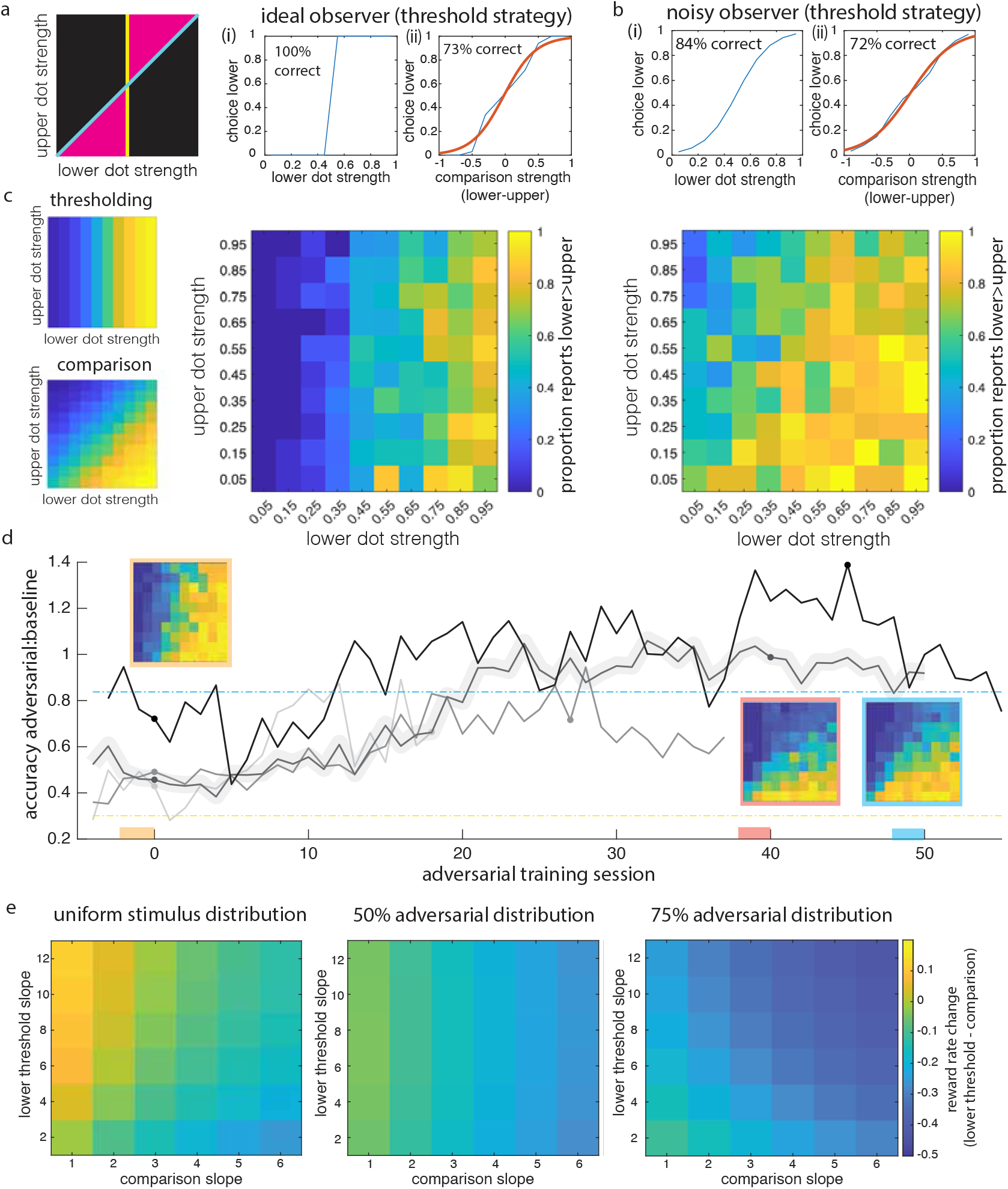
Changing stimulus distribution shifts behavioral strategy. a) Candidate decision strategies represented as boundaries in stimulus space. An ideal observer carrying out a single-region threshold strategy (e.g. yellow decision boundary) would achieve 100% accuracy (i) if that were the true task criterion (by definition), but only 75% accuracy on average and 73% accuracy on the specific simulation shown (ii) if the task required comparison between regions. b) The presence of sensory noise degrades the difference in reward return rate between the two strategies. c) 2D choice profiles allow readout of decision strategy. Left: 2D psychometric of simulated choices over 800 trials. Right: 2D psychometric for two example rats showing a bias toward one of the two strategies, pooled across the last 3 sessions prior to adversarial training (Methods). d) Adversarial training shifts behavioral strategy over sessions. Adversarial accuracy metric for simulated agents carrying out the comparison (blue dashed line) or thresholding (yellow dashed line) strategy is shown for a uniform stimulus distribution. Shaded trace shows strategy trajectory of an example rat, with summary psychometrics pooled over 3 sessions prior to adversarial training (orange), at the end of adversarial training (peach), and 8-10 sessions after the end of adversarial training (blue). Dot markers denote the start and end of adversarial training for each animal. e) Altered stimulus distribution in adversarial sessions shifts the reward rate cost of choosing the thresholding strategy over the comparison strategy.

In order to reliably read out strategy from choice profiles, we “unrolled” the psychometric function into the two-dimensional stimulus space from Figure 2a. By plotting the binned choice across this stimulus space, we could read out if subjects employed a thresholding (vertical or horizontal stripes) or comparison (diagonal stripes) strategy (Figure 2c). Again we found that across the 4 rats trained on this task, animals varied in the strategy they used to solve the task, but all rats fell somewhere between the two extremes (Figure 2c,d).

We wondered whether the strategy used by a given animal was fixed or if it would adapt to changing stimulus probabilities. Until now, we had only tested animals on a uniformly drawn distribution of stimuli, and we reasoned that changing the stimulus distribution to overemphasize the stimuli that distinguished between the two extreme strategies (magenta stimulus regions, Figure 2a) might push animals to adapt their strategy. We termed these stimuli “adversarial” stimuli, as they challenge the predominant strategy choice of the animals and distinguish between the target strategies. We then trained animals on sessions with increasing proportions of these trials, first by doubling the proportion of trials in this region then tripling it, a process we termed “adversarial training”. We found that animal strategy was indeed flexible to this manipulation of the stimulus distribution: within 10-20 sessions of adversarial training, all animals had shifted their strategy, as demonstrated by the increase in the ratio of accuracy on adversarial vs baseline stimuli (Figure 2d). Three of the four rats were returned to a uniform distribution of stimuli following adversarial training, and we found that their adversarial trial accuracy slowly decayed over the next 10 sessions, with varying speeds, further suggesting that animals adapt their decision strategy to recent statistics of the environment.

## Discussion

We demonstrated in this study that probe trials delivering stimuli that are adversarial to a given behavioral strategy can be used to distinguish between candidate strategies for solving a binary decision. We found that animals trained on the same task showed variation in the strategies they used to solve the task. Furthermore, we modulated the stimulus distribution to overrepresent the stimuli that were adversarial to the preferred strategy to investigate the effect on decision strategy selection. Using this approach, we found that in a visual decision task, rats largely preferred a single-region thresholding strategy over a two-region comparison strategy, but flexibly adjusted this strategy when exposed to extreme stimulus distributions that increased the reward cost of their preferred strategy. However, this cost had to be extreme to trigger a change in strategy, which was gradual and accumulated over days.

That animals sometimes behave suboptimally has also been found in other decision contexts, in particular when integrating across modalities (Carandini, 2024). However, our findings suggest that the form of suboptimality observed here is not an upper limit on animal behavior, but rather, can be modulated in response to context and task demands. Why might animals predominantly prefer a suboptimal strategy in the first place? In a previous study, we found that roughly equal numbers of primary visual cortex (V1) neurons preferentially fired to the two easiest visual stimuli (Zhang and Zador, 2023), so information about both lower and upper visual streams are available at least until V1. Further, the fact that animals can shift towards a comparison strategy when it is much more reward optimal suggests that there is no fundamental sensory processing limitation on comparison computations. However, one possibility is that thresholding operations are simpler or less effortful than comparison operations when stimuli vary across two dimensions (Scott et al, 2015, Akrami et al, 2018). A comparison operation may also accumulate more sensory noise, in which case the suboptimality of the strategy is offset by a higher sensitivity to the single stimulus stream the animal is using (Figure 2b,e).

Our results highlight the utility of probe trials in perceptual decision tasks to confirm the behavioral strategy used by individual animals. Our study leaves open the question of what are the neural correlates of a changing behavioral strategy, particularly when the strategy change involves a change in whether or how a portion of the stimulus is used. Future studies can begin to address these questions by recording neural activity during the adversarial learning process.

## Materials and Methods

### Ethics Statement

All procedures were approved by the Cold Spring Harbor Laboratory Institutional Animal Care and Use Committee (approval number 22-19-16-13-10-07-03-00-4), and conducted in accordance with National Institutes of Health guidelines.

### Animals and surgical procedures

Eight of nine rats (all five trained on the one-dimensional stimulus set, and three of four trained on the two-dimensional task variant) were male Long Evans rats obtained from Taconic Biosciences and Charles River at 8-10 weeks of age, and started training after reaching at least 10 weeks of age. The final rat expressed Cre-recombinase in all PV cells and was bred at Cold Spring Harbor Laboratory by crossing a PV-Cre+ BAC transgenic rat (Strain: LE-Tg(Pvalb-iCre)2Ottc, NIDA/NIMH) with a PV-Cre-rat. Rats were pair-housed for the duration of these experiments, in a reverse 12h light/dark cycle. Five of the nine rats were later implanted with microdrives and/or optical fibers for separate studies.

### Task design and behavioral system

The behavioral task has been previously reported (Zhang and Zador, 2023). Briefly, rats were placed in custom behavioral chambers containing three ports attached to a clear wall panel through which a monitor was visible. The Bpod system (Sanworks, NY) to implement the behavioral state machine and port entries and exits were detected using IR beam breaks. Animal entry into the center port triggered trial start and a pre-stimulus delay (duration drawn from an exponential function with a mean of 300ms). A visual stimulus was then delivered for 500ms using Psychtoolbox (Brainard, 1997; Pelli, 1997; Kleiner et al, 2007), followed by a 200ms fixed post-stimulus delay. Then, upon a decision tone, the animal was given 3s to make a decision by poking into a side port. A correct port nosepoke lasting at least 50ms triggered a 20 μL reward. No intertrial interval was specified following correct (either rewarded or missed reward) trials. A 1s punishment tone (white noise stimulus) and a 5-6s time out followed an incorrect choice. Aborting the trial at any point before the decision tone led to a missed trial condition, triggering a time-out of 2s.

All visual stimuli and auditory tones were delivered using Psychtoolbox. Visual stimuli were delivered at 60Hz refresh rate, with individual dots randomly distributed across each frame according to the stimulus condition on that given trial. The two stimulus subregions were of equal size, separated by a thin boundary region where no dots were ever present. Each dot location corresponded to a round white dot that subtended about 3^°^ in visual space. Dot locations were drawn from a uniform grid where every tenth pixel was a possible centroid. Of these possible locations, only 1% were selected as active on any given frame. Because dots were sparse, but dot size exceeded the spacing of the grid (30 pixel diameter), overlap was possible but minimal. Maximum overlap occurred on “95% probe” trials and trials in the upper right corner of the stimulus space (Figure 1) where the maximum number of dots were used in both subregions. In these trials overlap was on average 5% of the dot-occupied area, and did not exceed 11%.

In probe trials, the number of mean dots in each subregion was equalized. No correct response was specified for probe trials and side choices were rewarded randomly. In the 2D task, the number of dots shown in each region was independently drawn. To increase the proportion of “adversarial” stimuli during the “adversarial training” phase, all trials were first drawn from a uniform distribution, then trials with stimuli within the baseline region were selected at random to be redrawn within the adversarial stimulus region. The size of the biased draw subset is termed the “adversarial percentage”.

### Psychometric analyses

For the 1-D stimulus task (Figure 1a-c), choice proportions were calculated for each stimulus condition, and a logistic function was fit to the data:

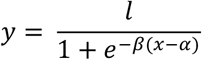

Where,

*y* is the choice proportion,

*x* is the stimulus condition,

*β* is the slope of the psychometric,

*α* is the intercept or bias, and

*l* is the maximum choice proportion (usually 1).

For the 2-D stimulus task (Figure 1d,e; Figure 2), stimulus strengths were discretized into 10 evenly spaced bins, and choice proportions were calculated for each corresponding bin. The same logistic function was then fit to this binned data.

### Psychometric simulations

To simulate choices made using either the thresholding or comparison strategy, 800 trials were drawn from a uniform stimulus distribution, and a logistic function as above was applied for each stimulus condition (as encoded by subregion strength or comparison strength) to generate a predicted choice proportion. The maximum choice proportion l was set to 1, the intercept was set to x = 0.5 for lower stimulus strength, or x = 0 for comparison strength (i.e., no bias), and the slope was set to 4 for the thresholding strategy (Figure 2b), which is similar to the logistic fits from the data, and varied for later analyses (Figure 2e). The psychometric slope selected for the comparison strategy was ½ of the slope for the thresholding strategy, following the reasoning that the comparison strategy would accumulate sensory noise from two stimulus subregions, whereas the thresholding strategy only required one.

To replot choice proportions from the thresholding strategy against the comparison strength of the stimulus, we assigned each trial a “choice” determined by the choice proportion for that given subregion strength, and then re-aggregated the trials according to the comparison strength, and summed the choice probabilities within these conditions. We then fit a psychometric function to these choice probabilities, as described above (Figure 2b(ii)).

### Adversarial trial accuracy metric

To summarize the strategy used within each session by each animal, we calculated a ratio of the accuracy on “adversarial” trials (magenta stimulus region in Figure 2a) to the accuracy on “baseline” trials (black stimulus region in Figure 2a). For an ideal observer, this would yield a ratio of 1 if the agent used the comparison strategy, and 0 if the agent used the thresholding strategy exclusively.

However, in the regime of sensory noise, and because these regions cover different distributions of comparison strengths, one would not expect an animal performing the comparison strategy to achieve an adversarial-to-baseline accuracy ratio of 1. As such, we computed this metric for the simulated choice profiles as described above, and confirmed that the comparison strategy returned values closer to 1 (0.8376 in the simulation shown here), whereas the thresholding strategy returned values below 0.5 (0.3010 in this particular simulation).

## Acknowledgements

This work was supported by grant R01DC012565 from the National Institutes of Health (A.M.Z).

## Declaration of Interests

A.M.Z. consults for and is a founder of Cajal Neuroscience.

## References

Akrami, A., Kopec, C. D., Diamond, M. E., & Brody, C. D. (2018). Posterior parietal cortex represents sensory history and mediates its effects on behaviour. Nature, 554(7692), 368–372. 10.1038/nature25510

Brainard, D. H. (1997). The Psychophysics Toolbox. Spatial Vision 10(4), 433–436. 10.1163/156856897X00357

Carandini M. (2024). Sensory choices as logistic classification. Neuron. 112(17):2854–2868.e1. 10.1016/j.neuron.2024.06.016.

Drieu, C., Zhu, Z., Wang, Z., Fuller, K., Wang, A., Elnozahy, S., & Kuchibhotla, K. (2025). Rapid emergence of latent knowledge in the sensory cortex drives learning. Nature, 641(8064), 960–970. 10.1038/s41586-025-08730-8

Kleiner, M., Brainard, D., Pelli, D., Ingling, A., Murray, R., & Broussard, C. (2007). What’s new in psychtoolbox-3. Perception, 36(14), 1–16.

Lak, A., Costa, G. M., Romberg, E., Koulakov, A. A., Mainen, Z. F., & Kepecs, A. (2014). Orbitofrontal cortex is required for optimal waiting based on decision confidence. Neuron, 84(1), 190–201. 10.1016/j.neuron.2014.08.039

Lee, K., Lee, K., Lee, H., Shin, J (2018). A Simple Unified Framework for Detecting Out-of-Distribution Samples and Adversarial Attacks. arXiv:1807.03888. 10.48550/arXiv.1807.03888

Pelli, D.G. (1997). The VideoToolbox software for visual psychophysics: Transforming numbers into movies. Spatial Vision 10(4):437–442. 10.1163/156856897X00366

Scott, B. B., Constantinople, C. M., Erlich, J. C., Tank, D. W., & Brody, C. D. (2015). Sources of noise during accumulation of evidence in unrestrained and voluntarily head-restrained rats. eLife, 4, e11308. 10.7554/eLife.11308

Waskom, M. L., Okazawa, G., & Kiani, R. (2019). Designing and Interpreting Psychophysical Investigations of Cognition. Neuron, 104(1), 100–112. 10.1016/j.neuron.2019.09.016

Zhang, A., & Zador, A. M. (2023). Neurons in the primary visual cortex of freely moving rats encode both sensory and non-sensory task variables. PLoS biology, 21(12), e3002384. 10.1371/journal.pbio.3002384

